# Insect asynchronous flight requires neural circuit de-synchronization by electrical synapses

**DOI:** 10.1101/2022.02.02.478622

**Authors:** Silvan Hürkey, Nelson Niemeyer, Jan-Hendrik Schleimer, Stefanie Ryglewski, Susanne Schreiber, Carsten Duch

## Abstract

Despite profound mechanistic insight into motor pattern generation, for asynchronous insect flight – the most prevalent form of flight employed by >600.000 species – architecture and function of the underlying central pattern generating (CPG) neural network remain elusive. Combining electro- and optophysiology, Drosophila genetics, and mathematical modelling, we uncover a miniaturized circuit solution of motoneurons interconnected by electrical synapses that, contrary to doctrine, serve to de-synchronize network activity. This minimal gap-junctional motoneuron network suffices to translate unpatterned premotor input into stereotyped firing sequences which are conserved across species and generate stable wingbeat power. Mechanistically, network de-synchronization requires weak electrical coupling in conjunction with specific postsynaptic excitability dynamics, revealing an unexpected, generic feature in the control of neural circuit dynamics by electrical synapses.

**One Sentence Summary:** Electrical synapses de-synchronize neural network firing to enable stable wingbeat power during insect flight.

## Main Text

With over a million known species, insects comprise the largest group of animals on earth (*1*). Their remarkable evolutionary success has been attributed to small body size and the ability to fly. These two features provide access to unutilized niches and rapid translocation, but aerodynamic constraints in small flyers require high wingbeat frequencies, and space constraints demand miniaturization of the central nervous controllers for flight (*2*). In 75% of all flying insect species, highly specialized asynchronous indirect flight muscles (a-IFMs) form an oscillatory system that generates wingbeat frequencies of 100-1000 Hz to ensure forward propulsion at low Reynolds numbers (*3*,*2*). This wingbeat oscillator is controlled by a central pattern generating (CPG) network in the central nervous system (CNS). Although asynchronous flight has emerged independently 7-10 times during evolution (*4*), neither the principles of CPG architecture for generating patterned motoneuron (MN) output from the miniaturized CNS of asynchronous flyers, nor the functional consequences thereof are known.

### Asynchronous flight is controlled by stereotyped motoneuron out-of-phase firing

To quantify asynchronous flight patterns and decipher CPG architecture we use the firing output of the five identified motoneurons, MN1-5 (*5*) innervating the dorsal longitudinal wing depressor muscle (DLM) of the genetic model system, *Drosophila melanogaster*, as well as other insect species to test for generality. The DLM provides the force for wing downstroke, consists of 6 muscle fibers, each of which is innervated by one identified MN (*6*,*7*) (Fig. 1B). MN1-4 each innervates one of the 4 most ventral ipsilateral DLM fibers, whereas MN5 is located contralaterally and innervates DLM fibers 5 and 6 (Fig. 1A). This neuromuscular architecture is conserved across all insect species examined so far.

**Fig. 1.**
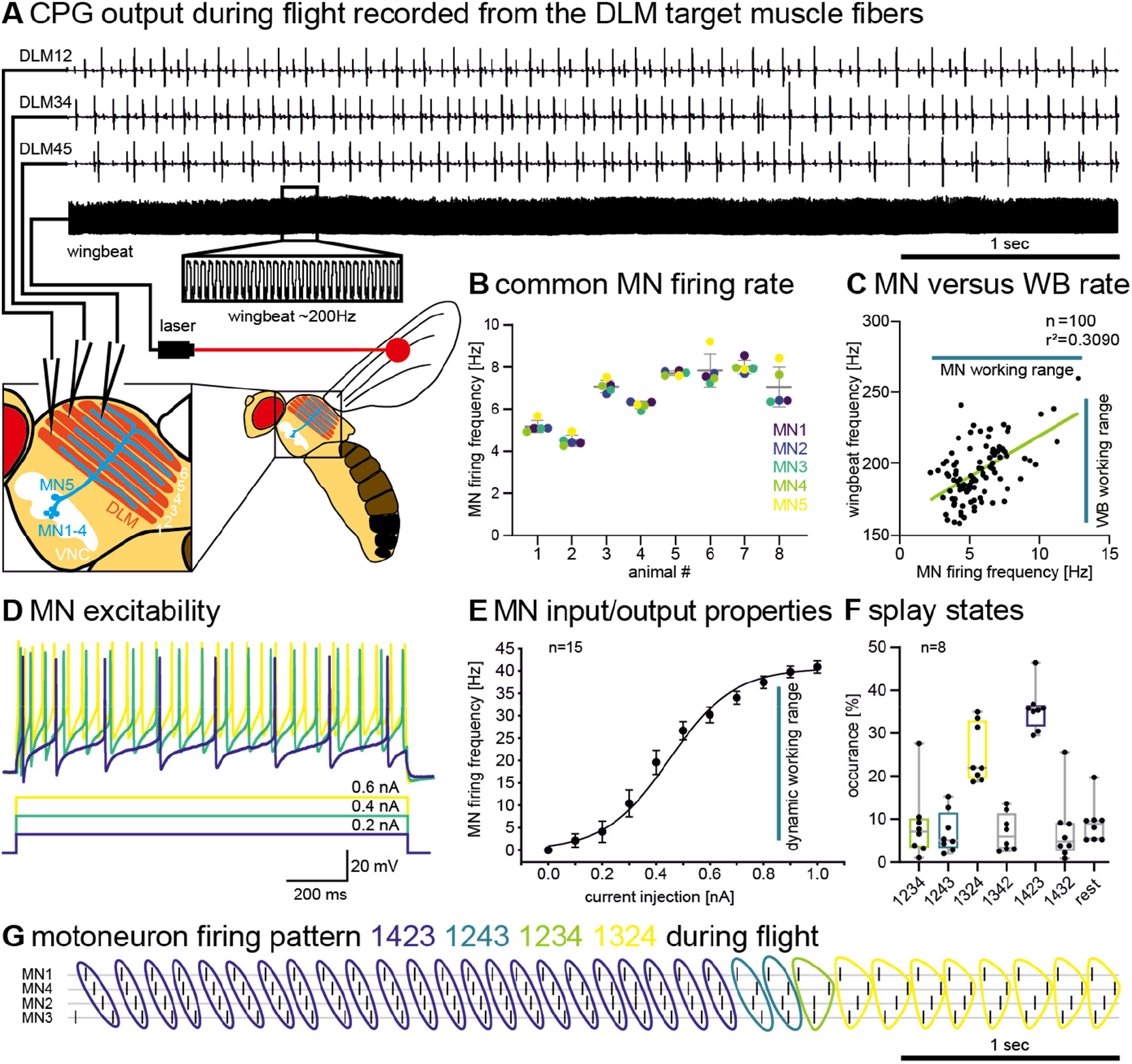
Centrally generated splay-state motor patterns control Drosophila flight. **(A)** Representative simultaneous recordings of all 5 DLM-MNs and wingbeat frequency (lower trace and black box for selective enlargement) during tethered flight. Schematic of DLM-MNs (MN1-5) in the ventral nerve cord (VNC) and their axonal projections to the 6 fibers of the DLM. **(B)** Firing frequencies of MN1-5 (color coded) are similar within a given animal. **(C)** MN firing- and wingbeat frequency (turquoise bars indicate normal working ranges) are related linearly proportionally (n = 100). **(D)** Representative tonic firing responses of DLM-MN to current injections. **(E)** Plotting injected current amplitude against response frequency (I/F) reveals a linear relation in the normal MN working range (turquoise bar) of 3-30 Hz (n=15). **(F)** During flight MN1-4 fire dispersed in time with characteristic sequences, which we name splay-states. Quantification of relative occurrence of different splay-states during 20 minutes of constant flight. Splay-states can change within one animal, but the same splay-states are preferred across individuals (n = 8). **(G)** Timing of MN1-4 spikes in four subsequent splay-states (1423, blue; 1243, turquoise; 1234, green; 1324, yellow) during flight.

*In vivo* recordings of MN1-5 from their DLM target muscle fibers with simultaneous laser-based wingbeat detection during tethered flight show that all five MNs fire only every ~40^th^ wingbeat (Fig. 1A-C). Although MN firing frequencies can vary between animals and are adjusted upon demand (*3*,*8*), within a given animal and power demand, all 5 DLM-MN always fire at the same frequencies (Fig. 1B).

MN1-5 firing frequencies directly control muscular tension and stretch activatability by adjusting myoplasmic calcium levels, and thus, wingbeat frequency and stroke amplitude (*3*,*9*). We confirmed with recordings from 100 animals that DLM-MN firing frequencies are directly proportional to wingbeat frequencies within the normal working range (Fig. 1C). Therefore, the CNS does neither control a-IFM power output by the recruitment of different motor units, nor on the scale of single wingbeats, but the frequencies of a-IFM-MN population firing are the key regulator of wing power production (*3*). We show that DLM-MN excitability is well adjusted to dynamically regulate wing power output: First, the MNs respond to input with slow tonic firing (Fig. 1D). Second, tonic firing responses exhibit nearly linear input-output computations at firing frequencies of 3-30 Hz (Fig. 1E), as occurring during flight (Fig. 1C). Therefore, MN excitability is perfectly adjusted to linearly translate synaptic input into wing power output in the working range of natural flight.

Despite equal firing frequencies at any given wingbeat frequency, (Figs. 1A, B) (*3*,*5*), the five DLM-MNs do not fire in synchrony. Instead, MN1-5 spikes are splayed-out in time with neuronal firing phases dispersed approximately equidistantly, which results in stereotyped preferred sequences of MN1-5 firing (Fig. 1G), as previously suggested (*5*). Importantly, the sequence of MN1-5 firing can change, but the CPG robustly slides back into one or two of the most preferred sequences, which we name the preferred splay-states. Strikingly, the same splay-states are preferred across animals (Fig. 1F), indicating that they result from hard-wired CPG circuitry. Splayed-out firing at preferred sequences results in characteristic phase relationships between any given pair of MNs, which again are conserved across individuals (Fig. S1). Characterization of phase relationships between all pairwise combinations of the five DLM-MNs allows testing for across species conservation of the splay-state motor pattern. The comparison of DLM-MN phase relationships reveals a striking similarity between *Drosophila melanogaster* and the green bottle fly, *Lucilia spec*. (Figs. S2A, B). Similar phase relationships have also been suggested for *Calliphora terrae-novae, Eucaliphora lilaea*, and *Musca domestica* (*10*). Moreover, using the MN4/MN5 pair (Fig. S2C) indicates conservation of CPG architecture between two Drosophila species (*D. melanogaster* and *D. hydei*), other dipteran genera (*Calliphora* and *Musca*), and to a certain degree even between *Diptera* (e.g. flies) and *Hymenoptera* (e.g. honey bee). Given that asynchronous flight evolved multiple times independently (*4*), splay-state firing has likely provided selective benefits over millions of years. To gain mechanistic and functional insight into the CPG network controlling asynchronous flight, we next analyzed the network principles underlying splay-state motor pattern generation and the resulting functional benefits for flight performance.

### Splay-state firing is produced by a minimal motoneuron network that is de-synchronized by electrical synapses

From invertebrates (*11*) to mammals (*12*) the timing and patterns of MN activation underlying locomotion typically relies on networks of premotor interneurons. In contrast, we find that splayed-out MN firing is generated by interactions between the MNs themselves. First, unpatterned optogenetic activation of excitatory, cholinergic input to DLM-MNs increases MN firing frequencies without changing the phase relations between MN pairs (Fig. S3). Second, optogenetic stimulation of the 5 DLM-MNs during flight increases their firing frequency, and thus wingbeat power, but it does not change phase relations between MNs (Fig. S4). Therefore, the timing and pattern of MN activation does not require patterned activity of interneurons, but the DLM-MN ensemble constitutes a minimal CPG.

One possible mechanism to transform common excitatory input to an ensemble of MNs into dispersed splay-state firing is lateral inhibition among MNs by chemical synapses (*5*). We rejected this possibility by combining targeted genetic manipulation of DLM-MNs with *in vivo* recordings during flight. Knock-down of receptors for inhibitory transmitters (GABA-ARs, GluCl) increases DLM-MN firing frequencies, but phase relationships are not affected (Fig. S5). Lateral inhibition via chemical synapses is, therefore, not required for pattern generation.

Another possibility to create a neural network with DLM-MNs only is to connect them with electrical synapses. This has previously been suggested (*13*,*14*), but experimental evidence has been lacking and electrical coupling seems difficult to reconcile with de-synchronized splay-state firing. We tested this by genetically manipulating innexins (15), the invertebrate counterparts of connexins (*16*) which comprise the pore forming proteins of electrical synapses. ShakingB (ShakB) is the innexin expressed in the Drosophila escape circuit, including the DLM-MNs (*17*).

Genetic manipulation of ShakB in DLM-MNs disrupts the splay-state without affecting firing frequencies (Figs. 2A, B). As exemplified for the MN4/MN5 pair, in control animals (n= 7) both MNs occasionally fire simultaneously (Fig 2A, red arrow), but firing of MN5 is inhibited shortly before and shortly after the occurrence of MN4 spikes. Consequently, phase histograms demonstrate preferred out-of-phase firing of MN4 and MN5 with a marked depression of MN5 that is centered around the MN4 spike (Figs. 2A, B, top). By contrast, with ShakB knock-down both MNs fire preferred in phase (Fig. 2A, B, middle). Therefore, electrical synapses are required for firing de-synchronization in this small CPG. Reducing the de-synchronizing function of electrical synapses by ShakB-kd in DLM-MNs impairs normal MN phase relationships and thus eliminates the preferred splay-state (Fig. S6, compare Fig. S1). This contradicts the common notion that electrical synapses synchronize network activity.

**Fig. 2.**
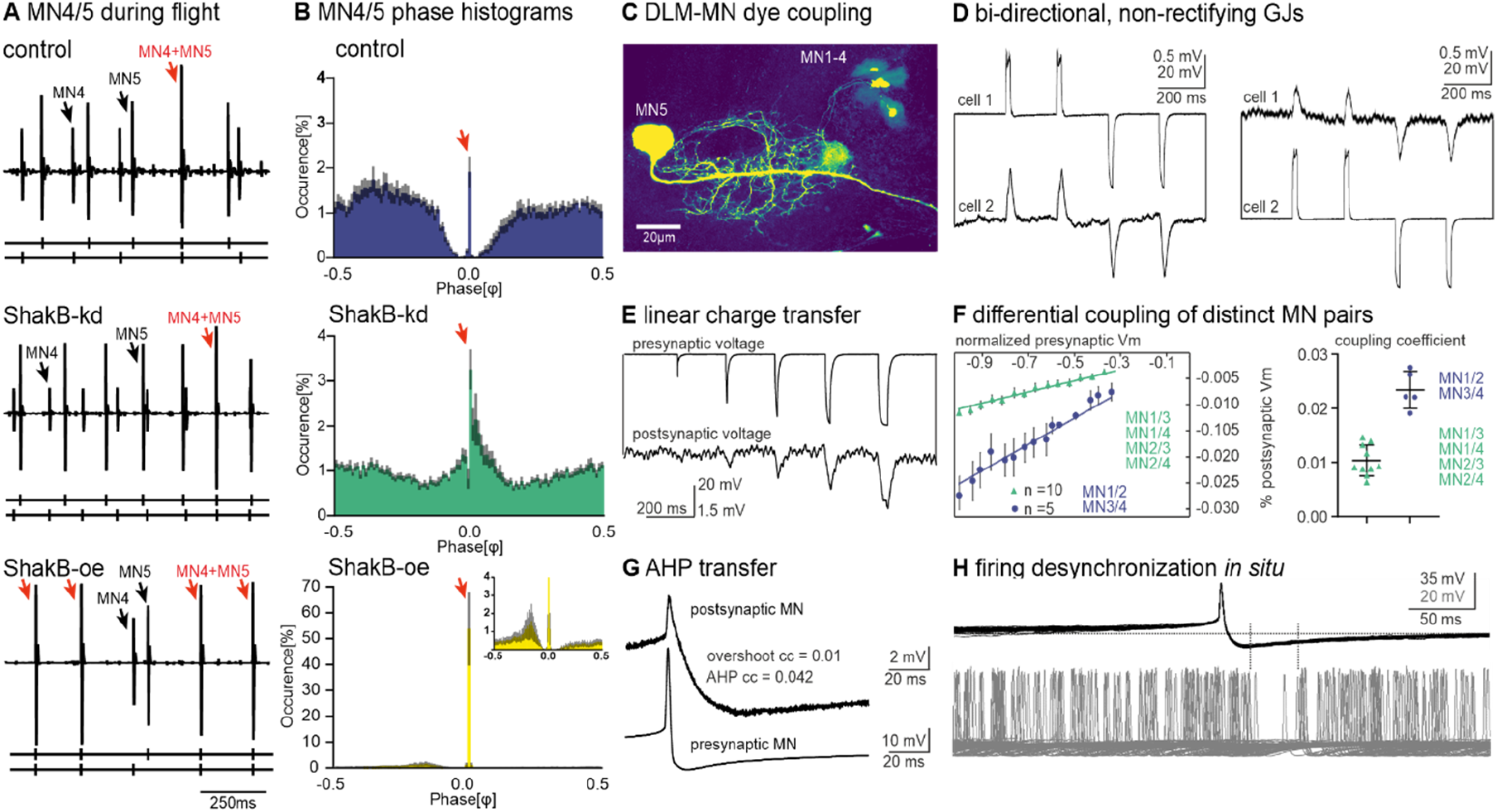
Electrical synapses shape CPG output by de-synchronizing MN firing. **(A)** Representative recording of MN4 and MN5 during tethered flight for control (top), RNAi-kd of ShakB encoded electrical synapses (middle), and overexpression of ShakB in MN1-5 (bottom). Black arrows demark MN4 and MN5 spikes, red arrows simultaneous MN4/MN5 spikes. **(B)** Phase histograms of occurrence of MN4 spikes (y-axis) in relation to consecutive MN5 spikes (phase 0 = MN5 spike) for control (top), ShakB-RNAi-kd (middle), and ShakB-overexpression (bottom). N=10 for each genotype. Red arrows demark synchronous MN4/MN5 firing. **(C)** MN1-5 dye coupling upon increasing GJ strength with dfmr1-RNAi-kd. **(D-H)** Intracellular recordings of MN pairs: **(D)** Hyper- and depolarizations are conducted bidirectionally (from cell 1 to cell 2 and *vice versa*). **(E)** Increasing current injection amplitude (upper trace) increases response amplitudes in electrically coupled MNs (lower trace). **(F)** Plotting pre-against postsynaptic voltage (left) reveals linear relations, but regression slopes differ between distinct MN pairs. Coupling coefficients (CCs, right) differ significantly between MN1/2, MN3/4 (blue) and all other pairs (green). **(G)** CC for the spike afterhyperpolarization (AHP) is higher than for the spike overshoot. **(H)** Firing of a coupled MN ceases during the presynaptic MN AHP.

Key to this unexpected role of gap junctions is weak electrical coupling. The theory of coupled phase oscillators (*18*) suggests that a strong coupling of neurons causes synchronization. Overexpression of ShakB in DLM-MNs, therefore, should cause firing synchronization, which is indeed what we find (Figs. 2A, B, bottom). The *in vivo* data indicate that weak electrical synapses cause firing de-synchronization, but stronger coupling synchronizes firing. Weak electrical coupling between DLM-MNs is supported by anatomical experiments. Although we have previously not observed diffusion of small dye tracer molecules through gap junctions between DLM-MNs (*8*,*19*), GJ coupling strength can be increased by knock-down of the fragile X mental retardation protein (FMRP)/dfmr1 (*20*). In this genetic background dye coupling of all DLM-MNs is reliably observed (Fig. 2C).

In sum, a minimal CPG of electrically coupled MNs is sufficient to produce splay-state firing across power demands and does not rely on additional interneurons or chemical synapses, because unpatterned optogenetic activation of either presynaptic cholinergic neurons or MN1-5 during tethered flight increases MN firing rates and wingbeat frequency (Figs. S3, S4), but the phase relations between MNs remain unaltered (compare Figs. 2A and S3B, S4B). But how do electrical synapses cause firing de-synchronization?

### Electrical synapses as required for motoneuron network de-synchronization are weak, bidirectional, and non-rectifying

Dual *in situ* patch-clamp recordings of MN pairs confirm electrical coupling and characterize the electrical synapses between MNs as weak, bidirectional, and non-rectifying (Fig. 2D). Non-rectifying because depolarizing or hyperpolarizing current injection (20ms duration) into one MN causes gap junctional (GJ) potentials in the other MN. The relationship between pre- and postsynaptic charge is linear (Figs. 2E, F). Bidirectional because the direction of charge transfer can be reversed (Fig. 2D). Compared to common coupling coefficients (CCs, postsynaptic charge/presynaptic charge) as estimated in different types of neurons (0.02-0.2;(*21*)) electrical synapses between DLM-MNs are weak, but coupling is twice as strong for the MN1/MN2 and MN3/MN4 pairs (CC=0.023±0.003) as compared to all other possible combinations of MN1-4 pairs (CC=0.01±0.0027, Fig. 2F). Weak electrical synapses between all MNs, but different CCs between distinct pairs of MNs is key to generating the splay-state (see below).

The possibility that electrical synapses do not always synchronize the activity of coupled neurons, but may de-synchronize neural networks under specific conditions has been suggested by theoretical work (*22*–*24*). Experimentally, a transient de-synchronization of electrically coupled cerebellar Golgi Cells (GCs) has been described in one study, but for the specific condition of sparse input (*25*). There, transient firing de-synchronization has been attributed to a more effective gap junctional transmission of the action potential (AP) afterhyperpolarization (AHP) as compared to the depolarizing AP overshoot (*25*). However, the pre-conditions for desynchronization of electrically coupled cerebellar GCs are not fulfilled in the insect asynchronous flight CPG. First, during flight network de-synchronization by electrical synapses is not transient but permanent. Second, network de-synchronization occurs not only under sparse synaptic input regimes, but through the full range of synaptic input regimes that can occur during flight. Therefore, additional mechanisms for gap junction mediated firing de-synchronization are likely required for splay-state generation.

However, we detect differentially effective transmission of different parts of the presynaptic spike through electrical synapses. In fact, the AHP of the DLM-MN spike is transmitted more effectively through ShakB mediated electrical synapses and shows a significantly higher CC (0.042±0.04) than the brief spike overshoot (CC, 0.01±0.004; Fig. 2G). CCs can differ for different components of the AP, because the duration of the presynaptic signal and the time constant of the postsynaptic membrane shape the junction potential (*21*). Paired *in situ* current clamp recording of MNs, that were induced to fire tonically at 2-10 Hz by somatic current injection, indicate that firing of one MN can depress firing of the other one during and shortly after the AHP (Fig. 2H). This is in agreement with the findings on transient de-synchronization of GC firing in cerebellum (*25*). However, it seems unlikely that this mechanism suffices for firing de-synchronization in the flight CPG. First, AHP mediated DLM-MN firing suppression (Fig. 2H) is not robust across different MN firing frequencies as occurring during flight. Second, MN firing is not only robustly de-synchronized across firing rates and power demands, but it is also organized into specific firing sequences, splay-states (Fig. 1F). Therefore, splay-state generation likely requires additional or other mechanisms. To address these, we turn to theoretical considerations.

### Network de-synchronization requires weak electrical coupling and a specific neuron-type

What are possible general mechanisms for small network de-synchronization without a pronounced AHP (likely not sufficient, see above) or inhibitory synapses (not required for DLM-MN de-synchronization, Fig. S5)? According to the theory of phase oscillators, networks can display a splayed-out state if pairs of neurons have a preference to fire out-of-phase, i.e., they are phase-repulsive (*18*). For networks of more than two neurons, the antiphase state, however, cannot be achieved by all neuronal pairs; these systems are called frustrated (*26*,*27*) and can organize into a splay-state, which equidistantly maximizes phase distances among neurons to minimize frustration. In neurons, phase-repulsive coupling is fostered by a specific excitability class, the homoclinic orbit (HOM) spike-onset (*28,18,29*), which is shaped by cellular properties, including their ion channel composition. To test the hypothesis that weak electrical synapses between neurons with HOM bifurcation provides a mechanism for firing de-synchronization in small neuronal networks, we used a minimal two conductance-based model fitted to DLM-MN sodium and delayed rectifier ion channel kinetics (*30*; suppl. materials) to generate a gap junctional network of MN1-4 (Fig. 3A). Subjecting the single neuron model to a bifurcation analysis reveals the presence of an excitability switch (*29*), with a HOM spike-onset for moderate expression values of Shab delayed rectifier channels (Fig. S7A and suppl. materials). Networks of four identically, weakly electrically coupled DLM-MNs with HOM excitability robustly exhibit de-synchronized firing (Fig. 3B). Note that the minimal conductance-based model does not contain a pronounced AHP (see spike shape in Fig. 3B), indicating that the presynaptic AHP is not mandatory for network de-synchronization. By contrast, sensitivity analysis reveals that weak electrical coupling is required, because all models with small CCs (< 0.2), as obtained *in vivo* (Fig. 2F), yield de-synchronized firing (Fig. 3C), whereas with CCs > 0.2 network synchronization increases (Fig. 3C). Moreover, transforming postsynaptic neuron excitability from HOM to different bifurcation classes (SNIC in Fig. 3D) by increasing Shab delayed rectifier channel levels also causes firing synchronization and destroys the splay-state. Therefore, weak electrical coupling between homoclinic neuron types provides a general mechanism that is sufficient to produce firing de-synchronization in small networks. Importantly, this mechanism holds across different firing rates and does not require a pronounced AHP, although the AHP may further stabilize de-synchronization.

**Fig. 3.**
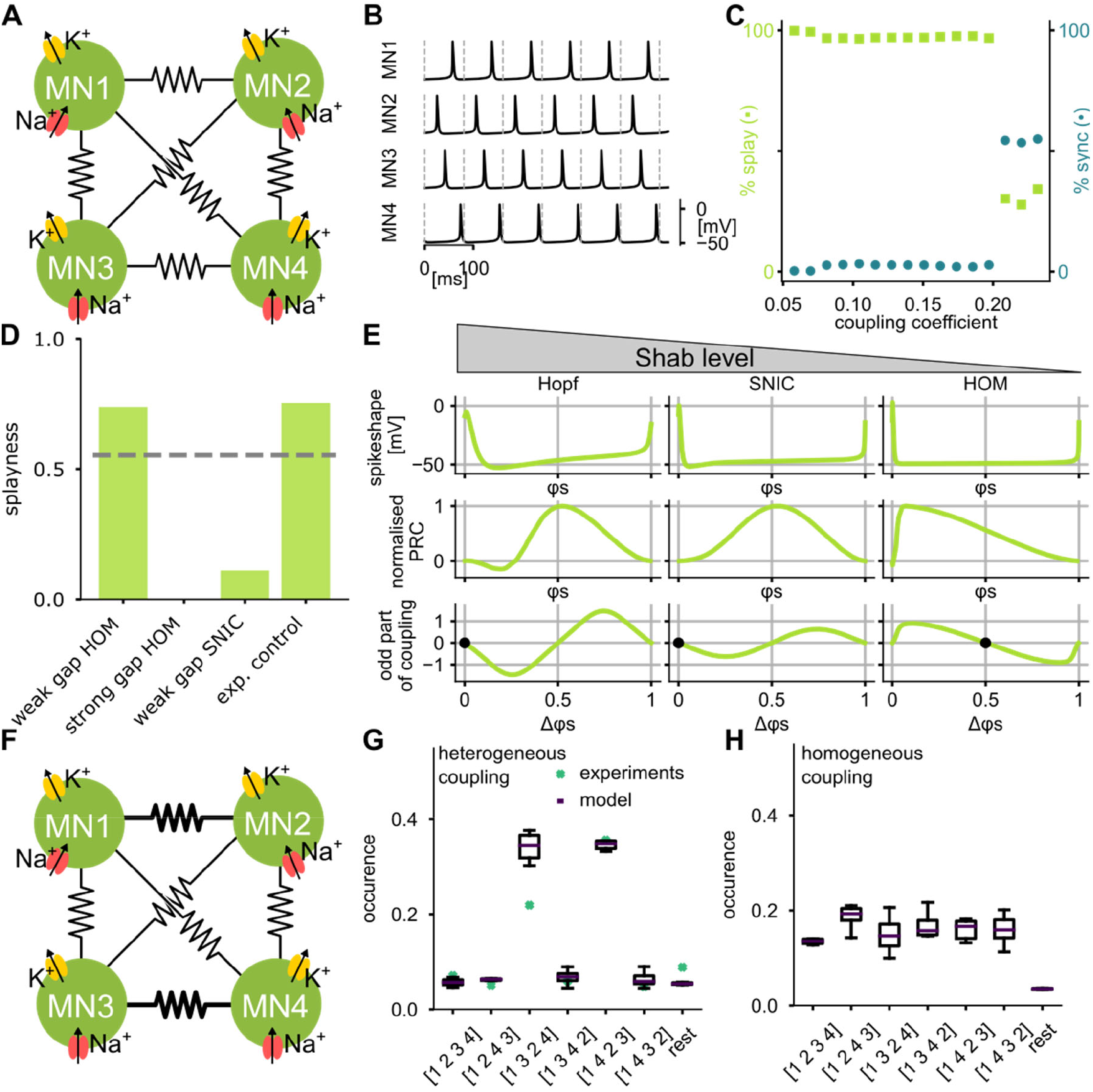
Weak electrical synapses and a HOM PRC underlie firing de-synchronization in small gap junctional networks. **(A)** Schematic of conductance-based model with identical weak GJs between all MNs. **(B)** Network produces splay-state firing **(C)** with electrical CCs <0.2. Percentage of in-phase synchronized networks increases at CCs >0.2. Gap junction effects on intrinsic neuron properties were compensated and states were quantified using order parameters (suppl material). **(D)** Splay-state requires weak coupling and a HOM PRC. Splayness index (suppl material equation 7) shown for 600s of simulation with noise per condition. Gray line represents random phase relations. **(E)** Increasing Shab conductance levels transforms the MN PRC from HOM via SNIC to Hopf types. Different spike-onsets vary in AP waveform (top row) and PRC (middle row). Averaging theory (suppl material equation 4) yields the odd part of the coupling function for each phase distance Δφ, which represents the effective phase shifts during tonic firing (bottom panel). Only a HOM PRC yields one stable fixpoint at phase 0.5, thus favoring anti-phase firing. **(F)** Adding noise and different CCs for different MN pairs to the model produces the same preferred splay-states as observed *in vivo* **(G)**, while homogenous coupling fails to do so **(H)**.

### Mechanism of de-synchronized splay-state firing in a frustrated network

Our theoretical analysis suggests that the effect of synaptic transmission through the same gap junction can be reversed by the expression of different ion channels in the electrically coupled neuron, thus resulting in either network synchronization or de-synchronization (Fig. 3D). Due to the generic nature of the mechanism, analytically derived from the theory of coupled phase oscillators, the unanticipated de-synchronizing role of electrical synapses for neural network dynamics is a universal feature of conductance-based models with HOM phase response curves (PRCs). Specifically, given that synaptic input drives DLM-MNs in a tonic spiking regime (Fig. 1A), it is possible to reduce the conductance-based model to approximate phase oscillators (*31*) defined by their intrinsic mean firing rate and PRC (see suppl. materials). Analysis of a reduced phase response model confirms de-synchronization of weakly coupled neurons with an asymmetric PRC, such as a HOM spike-onset (Fig. S7). Moreover, PRC analysis reveals that electrical coupling of homoclinic MNs yields a frustrated system whose solution includes the splay-state. For all bifurcation classes that can be reached by tuning Shab channel levels, the postsynaptic voltage perturbation is calculated between a pair of pre- and postsynaptic firing phases. The perturbations depend on the spike waveform (Fig. 3E, top row), which differs among excitability classes, and the GJ properties (non-rectifying, bidirectional as measured *in vivo*, Figs. 2D-F). To map the perturbations to shifts in spike timing of the coupled neuron, the PRCs of the excitability classes are used (Fig. 3E, middle row). For reciprocal coupling, the phase relations of tonically firing neurons can be read from the odd part of the coupling function (supplementary material, Eq. (3)) and critically depend on the PRC shape (Fig. 3E, bottom row), which in turn depends on the bifurcation type of the neuron (Fig. S7A). For example, for a SNIC PRC a small phase distance, Δφ, (neuron 1 fires shortly before neuron 2) is further decreased, whereas a large Δφ (neuron 2 fires shortly before neuron 1) is further increased. This results in a stable fixpoint at phase 0, so that the neurons show synchronized firing (Fig. 3E, bottom). In contrast, with a HOM PRC a small Δφ causes neuron 1 firing to be accelerated, whereas a large Δφ causes neuron 2 firing to be accelerated. This results in a stable fixpoint at Δφ=0.5 for each coupled neuron pair. For a network of >2 electrically coupled neurons this causes a frustrated state, because antiphase locking at Δφ =0.5 cannot be achieved for all units at the same time. In a small network of 4 to 5 neurons, such as the flight CPG network, the splay-state represents a low-frustration solution which determines the most likely network state (*32*).

### Sequence preference requires heterogeneity in coupling

Our *in vivo* recordings show that DLM-MN firing is not only de-synchronized, but in addition organized into preferred sequences of splayed-out firing, which are conserved across individuals (Figs. 1F, S1) and species (Fig. S2). How are the preferred sequences generated? Adding noise and heterogenous electrical coupling strength (Fig. 3F) as observed *in situ* (Fig. 2F) produces the same preferred splay-states with similar sequence statistics (Fig. 3G) as observed *in vivo*. By contrast, a network with homogenous electrical coupling fails to show sequence preference (Fig. 3H). We conclude that heterogenous weak electrical coupling of homoclinic neuron types sufficiently explains the firing sequences of the preferred splay-states.

### Splay-state firing is required for stable wing power production

The conservation of the preferred splay-states of DLM-MN firing across individuals and species suggests that MN firing de-synchronization provides a significant functional benefit. This can be tested by increasing electrical coupling via genetic manipulation to transform splayed-out into synchronized MN firing (Fig. 2A and 3C). *In vivo*, MN firing synchronization induces fluctuations in wingbeat frequency (Fig. 4A). Each synchronized MN spike is followed by a peak in wingbeat frequency (Figs. 4A, yellow) with a latency of 7.3 ± 1.3 ms (Fig. 4C). By contrast, wingbeat frequency fluctuations during splay-state firing are 4 to 5-fold smaller and not correlated in time with the spikes of any of the DLM-MNs (Figs. 4A, C, blue).

**Fig. 4.**
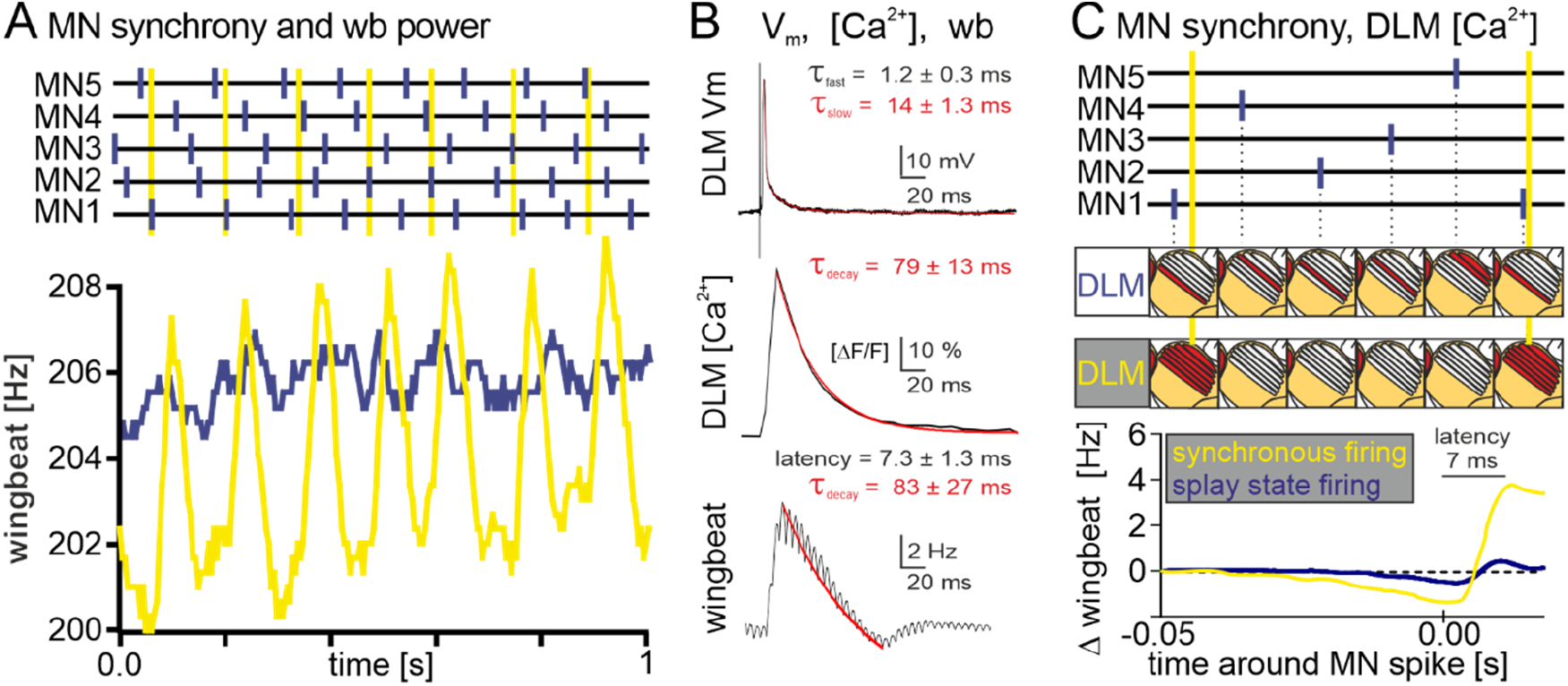
Splay-state firing ensures stable wingbeat power. **(A)** Splay-state firing in controls (top panel, blue) as compared to synchronized firing as observed with overexpression of ShakB in MN1-5 (top panel, yellow). Splay-state firing results in constant wingbeat frequency (lower panel, blue), but synchronized firing in fluctuating wingbeat frequencies over time. **(B)** Representative intracellular current clamp recording from DLM fiber 5 (inset shows magnification) in response to MN5 firing (top). Middle panel shows DLM fiber calcium signal (ΔF/F) imaged with expression of GCaMP8f in response to a single DLM fiber spike (red line is decay fit). Lower panel shows the time course of wingbeat frequency changes after synchronous MN1-5 spikes. Latency depicts the delay between MN spiking and the peak wingbeat frequency (~7 ms). **(C)** Splay-state firing (blue) causes alternating calcium signals (depicted as red DLM fibers schematics) in the 6 DLM muscle fibers, whereas and synchronized firing (yellow) causes synchronized calcium signals across all 6 DLM fibers. Timing of average changes in wingbeat frequency in relation to synchronized MN spiking (yellow, n = 3) and to MN firing in the splay-state (blue, n = 3).

Under steady state conditions, MN firing rates are directly proportional to myoplasmic calcium levels and wingbeat power (*3*), but the dynamics of myoplasmic calcium and power output after single MN spikes are not known. Therefore, we measured the temporal relationships between muscle fiber depolarization, changes in myoplasmic calcium levels, and power output (Fig. 4B). DLM muscle fiber APs are carried by sodium and calcium and exhibit two time constants of voltage decay, with τ_slow_ ~14 ms, likely resembling the calcium-carried component (Fig. 4B, top). In response to MN firing, myoplasmic calcium levels as monitored *in vivo* with GCaMP8f expression in flight muscle peak within 10 ms after the MN spike and show a decay τ of ~79 ms (Fig. 4B, middle). Strikingly, after an AP-evoked rise in myoplasmic calcium wingbeat frequency increases quickly within 7.3 ms (Figs. 4B, C) and decays with a τ of ~83 ms. Therefore, the dynamic changes in intramuscular calcium as induced by single MN spikes match the dynamic changes in wingbeat power, with on-kinetics within 10 ms and off-kinetics within 80-100 ms. Moreover, the time course of myoplasmic calcium signal decay explains why splay-state firing causes homogenous wingbeat power over time, whereas synchronous firing causes power fluctuations, which result in stuttering flight patterns. Given that each of the six muscle fibers is innervated by one DLM-MN, even temporal dispersal of MN spikes will also splay-out the calcium fluctuations between the six muscle fibers (Fig. 4C). In the working range of normal flight (3-25 Hz MN firing) the myoplasmic calcium levels will decay in each DLM fiber in between two subsequent MN spikes to this fiber. Therefore, each fiber will exhibit fluctuations in stretch activatability over time. Splay-state firing ensures that the average power production across all 6 fibers is constant over time, because each fiber will show maximal stretch activatability at a different time point (Fig. 4C, top). By contrast, synchronous spikes of all DLM-MNs will cause synchronous calcium elevations in all 6 muscle fibers, which will result in common changes in myoplasmic calcium between two subsequent synchronous spikes, and thus in marked increases in wingbeat power within 7 ms after each synchronous spike followed by declining wingbeat power with a decay τ of ~80 until the next synchronous MN spike will occur (Fig. 4C, middle). Quantification of wingbeat frequency changes in temporal relation to MN spiking shows that synchronous firing (n=7) increases the amplitude of power fluctuations by a factor of ~8 in comparison to splay-state firing.

In sum, splay-state asynchronous flight motor patterns are conserved across individuals and species, are produced by a minimal CPG of weakly electrically coupled motoneurons, and serve constant wingbeat power output at a given power demand. This provides a comprehensive view of the asynchronous flight CPG network structure and the resulting functional consequences for the most abundant form of locomotion on earth. In addition, we provide a theoretical background for firing de-synchronization/synchronization in small gap-junctional networks. The underlying mechanism is generic, predicting de-synchronizing functions of electrical synapses beyond the motor system of insects. Electrical synapses can thus be employed for operations like sign-reversal and ensemble firing de-synchronization, a functional versatility and impact on neural circuit dynamics that was previously attributed to chemical synapses alone.

## Supporting information

Movie S1 Drosophila melanogaster

Movie S2 Lucilia spec

Movie S3 Apis mellifera

## Acknowledgments

We thank Olaf Buda and Andrij Bujnenko for help with tethered flight recordings as well as Profs. Drs. P.R. Hiesinger (Berlin), U. Thomas (Magdeburg), and M. Silies (Mainz) for many helpful comments on the ms.

## Author contributions

Conceptualization by all authors, investigation were conducted by SH, SR, NN, JHS; Formal analysis was conducted by SH, SR, CD, NN, and JHS. Data curation by NN, SH, SR. Project administration was led and Resources were provided by CD and SS. Validation and visualization by all authors. The ms was drafted by SH and CD and reviewed and edited by all authors.

## Data and materials availability

All data is available in the main text or the supplementary materials. All data, code, and materials used in the analysis will be available upon request.

