## Supplementary Materials for "Insect asynchronous flight requires neural circuit de-synchronization by electrical synapses"

**This PDF file includes:**

Materials and Methods
Supplementary Text
Figs. S1 to S7
Tables S1 to S4
Captions for Movies S1 to S3

**Other Supplementary Materials for this manuscript include the following:**

Movies S1 to S3

**Materials and Methods**

Animals

*Drosophila melanogaster* were reared in 68 ml transparent food vials (Kisker Biotech GmbH & Co. KG, 25mm x 95mm) filled with 10 ml of standard cornmeal/glucose/yeast/agar diet at 25°C and 60% humidity under a 12h light/dark cycle. For experiments adult 2-5 days old male flies were used. We used Canton-Special (Canton-S) as wildtype control. To selectively target the DLM-MNs with UAS-driven transgenes we used a split GAL4 driver line that combines GMR23H06 (BDSC# 49050 discontinued) and GMR30A07 (BDSC# 49512) from the Rubin Collection (*33*). The GMR23H06 enhancer expresses the activating domain (AD) and the enhancer GMR30A07 expresses the DNA-binding domain (DBD). Overlap of both expression patterns, and thus functional GAL4 expression is restricted to almost exclusively to the DLM-MNs. The following transgenes were expressed in MN1-5 to manipulate gap junction expression and function. ShakB RNAi (BDSC# 57706) targets all shakB isoforms and was used to knockdown gap junctions in DLM-MNs. To selectively knock out gap junctions in DLM-MNs we combined GAL4-mediated, tissue-specific expression of UAS Cas9 (BDSC# 54595) with UAS-TRiP-CRISPR knock out for ShakB (BDSC# 78593). For gain of function via overexpression of gap junction protein the shakB(N+16) isoform was used (kind gift of P. Phelan, University of Kent). Another reported means (*20*) to increase GJ coupling strength between identified adult Drosophila neurons is targeted expression of *FMRP$^{RNAi}$* (*w;UAS-FMRP$^{RNAi}$/23H06-ADZ UAS-CD4-td-GFP;30A07-DBD/+*). We used this approach only to enhance dye coupling following intracellular fills (see below), but not for physiological analyses because of the risk of unwanted side-effects of FMRP knock down. Knockdown of inhibitory synaptic transmission through glutamate-gated chloride channels (GluCls), or RDL GABARs was conducted by targeting the expression of UAS-RNAi$^{KK}$ for GluClα (VDRC# 105754) or UAS-Rdl-RNAi (VDRC#41103) to DLM-MNs with the DLM-MN GAL4 driver GMR23H06. To test for electrical synapse function by genetic manipulation, UAS-RNAi mediated knock down of ShakB was compared with UAS-ShakB overexpression and empty UAS-control with at least 7 biological replicates in each group. To test for effects of GABAARs and GluClRs on flight patterns, UAS-RNAi knock downs for each receptor type was expressed in DLM-MNs and the effect on flight patterns compared to the respective genetic controls, with at least 7 biological replicates in each group. For all genetic manipulations the order of experiments was fully randomized, and analysis was conducted blindly without knowledge of the genotype. All animals that were successfully recorded for at least 10 minutes of flight were included in the analysis. For *in vivo* optogenetic manipulation of the activity of DLMs MNs during flight we expressed UAS-XXL Channelrhodopsin (XXL-ChR) (BDSC# 58374; (*34*)) either under the control of our DLM-MN-specific split GAL4 driver (split GMR23H06+GMR30A07) to enable light activation of the MNs, or alternatively, in presynaptic cholinergic neurons under the control of Cha-GAL4 (*35*). Direct light activation of DLM-MNs during flight was used to test whether the central pattern generator (CPG) correctly shapes MN firing into preferred sequences even when firing rates are artificially increased. Light activation of presynaptic cholinergic neurons during flight was used to test whether the CPG translates artificially presented, unpatterned synaptic input into patterned output from MNs. For *in vivo* calcium imaging in DLM muscle during flight the genetically encoded calcium indicator UAS-GCaMP8f (*36*) was expressed under the control of Act88-GAL4 (*37*). Other insect species were obtained from pet shops as feed insects and *Apis mellifera* from the local apiculture of the Johannes Gutenberg - University of Mainz.

TRiP CRISPR/Cas9 knock-out

To selectively knock out gap junctions in DLM-MNs we combined GAL4-mediated, tissue-specific expression of UAS Cas9 with TRiP-CRISPR knock out for ShakB (*38,39*) from the Transgenic RNAi Project (TRiP,(*40*)).

The single gRNA from the targeted gene is ubiquitously expressed by the U6:3 promoter. The tissue-specific expressed Cas9 is then guided to the target coding sequence, where the gene is cleaved. The following Non-Homologous End Joining (NHEJ) is completing the somatic gene mutagenesis.

Electromyography/ extracellular recording of motoneurons

The *in vivo* activity of the DLM-MNs during tethered flight was monitored by extracellular voltage recordings from their respective muscle fiber. For Flies were briefly cold anesthetized (20 s in an empty 68ml plastic vial on ice), transferred dorsal side up onto a cold metal plate (~3°C), and glued (clear glass adhesive (Duro; Pacer Technology, Rancho Cucamonga, CA)) with head and thorax to a triangle-shaped tungsten hook (0.1 mm diameter). Curing of the glass adhesive was induced by exposure to UV light (Mega Physik Dental Cromalux-E Halogen Curing Light Unit) for 45 s. The flies were kept individually for 10 min to recover from the cold anesthesia. The rested flies were then mounted in the setup to a clamp attached to a micromanipulator. Tarsal contact to a small polystyrene bead prevented unwanted engagement into flight. Next a light barrier consisting of a red laser, an aperture to reduce beam width to appr. 1 mm, and a LED light sensor was positioned so that the light beam was broken by each wingbeat. After positioning of this laser-based wingbeat detector tungsten electrodes for extracellular MN recordings were inserted. A reference electrode was inserted into the last abdominal segment. To record the activity of MN5 and MN4 from the dorsal most DLM fibers we inserted one sharpened tungsten wire into the dorsal thorax in front of the anterior dorsocentral bristle, making sure not to cross the midline of the thorax. This resulted in extracellular recordings of two units that could easily be distinguished by amplitude and shape. Confirmation of unit identity was achieved by adjusting the tungsten insertion depths and monitoring the resulting amplitude changes of the recorded units. Since MN5 innervated the dorsalmost DLM fibers it is the first unit to appear upon shallow dorsal electrode insertion. Moving deeper lets a second unit appear that belongs to MN4 which innervated the fiber just beneath. This MN4 unit increases in amplitude with increased insertion depth in DLM4, while the amplitude of the MN5 decreases. This procedure allowed non-ambiguous allocation of the two units to MN5 and MN4. To record up to 5 DLM units simultaneously, additional tungsten electrodes were inserted. A second electrode four small bristles in front of the first electrode, which recorded MN4 and MN3. A third electrode was inserted anterior on the same line as electrodes 1-2 at the position where small bristles begin to appear on the thorax. Again, unit identification was conducted by altering electrode insertion depth at the beginning of each experiment. Starting from dorsal and slowly moving ventrally the first unit to appear is MN5, the second one MN4, the third one MN3 and so on. Electrode depths were adjusted until the subsequent unit that appeared while moving deeper had approximately half of the amplitude of the first unit, so that both were easily distinguishable. In some recordings it was also possible to pick up three clearly separable units simultaneously, but spike sorting was more time consuming. All extracellular recordings were amplified at 1000x with an AM-Systems 1700 extracellular amplifier. Electrophysiological and wingbeat recordings as well as a frame trigger signal from high-speed video (see below) were digitized with an Axon Digidata 1550B (Molecular Devices),

acquired with AxoScope (Version 10.7) and stored on a PC. Spike sorting was conducted offline with the spike sorting function in Spike2 (Version 7.2). Data were further analyzed with Spike2, and custom written Python routines created with Jupyter notebook to count pattern probabilities and create phase histograms. For additional functions the Python libraries NumPy, pickle, SciPy, Matplotlib and seaborn were imported. All python script is available upon request. Additional data analysis and statistics were conducted with Microsoft Excel Professional Plus 2019 (Version 2110) and GraphPad Prism (Version 9.2.0).
All larger insect species than *Drosophila* were cold anesthetized for up to 20 min in the fridge at 5°C and fixated in a 3D printed device to hold the insect in place. The .stl file is available upon request. Instead of a rectangular tungsten hook, the wire is bent in a half circle and glued on the outer circumference of the posterior dorsal region of the thorax to increase the adhesive surface.

High speed video

In addition to electrophysiological recordings of DLM-MN activity and laser-based wing detection, selected times of tethered flight were simultaneously recorded as high-speed video with a Photron FASTCAM Mini UX100 and Photron FASTCAM Viewer software (PFV Version 3.6.9.0) at 5000 frames per s. As illumination two IR lights (Sygonix IR illuminator with 48 LEDs) were used. See supplementary materials for movies of Drosophila (movie S1), goldfly (movie S2) and honey bee (movie S3) tethered flight.

Electrolytic sharpening of tungsten electrodes

The tips of tungsten rods (diameter 100µm for Drosophila, 125µm for all larger insects) were sharpened electrolytically by repeated dipping in a $NaNO_2$ KOH solution (10.3 M $NaNO_2$ and 6.05 M KOH in dd$H_2O$.). Tungsten rods of 1 cm length were crimped onto metal rods taken from a circuit board connector. For this the wire was placed onto the rod and covered with a 0.5 mm ferrule. A crimping tool compressed it to a pluggable tungsten wire electrode. Now the electrode was placed into an alligator clip, connected to a stimulator (Grass SD9 square pulse stimulator) and repeatedly dipped into the sharpening solution, which was connected to the other pole of the stimulator. A monophasic current with 100Hz, 40V and 1ms duration was applied. The tip was frequently moved in and out of the solution to form a thin tip. Finally, electrodes were rinsed with distilled water.

Double Patch Clamp Recordings of DLM-MNs

After removing legs and wings, 2-3 days old adult Drosophila males were pinned into a Sylgard-coated lid of a 35 mm Petri dish and fixed ventral side up with minute pins in the head and through the tip of the abdomen. After submerging in normal saline (in mM: NaCl 128, KCl 2, $CaCl_2$ 1.8, $MgCl_2$ 4, HEPES 5, Sucrose ~35.5, pH was adjusted to 7.24 with 1 N NaOH, and osmolality was 300 mOsM/kg) the thoracical cuticle was removed with fine iris scissors to expose the ventral nerve cord (VNC). The specimen was then rinsed thoroughly with saline to remove excess debris. After mounting the preparation onto the stage of an upright Zeiss Axio Examiner epifluorescence microscope, the VNC was viewed with a 40x water immersion lens (Zeiss W Plan Apochromat 40x NA 1.0, DIC VIS-R), the UAS-6xmCherry-expressing DLM-MN1-4 (genotype: w;23H06-ADZ UAS-6xmCherry/+;30A07-DBD/+) were viewed through a TRITC filter set. The ganglionic sheath and debris hampering access to MN1-4 somata were removed by repeated application of 1% protease type XIV (from *Streptomyces griseus*, Sigma Aldrich, Cat# P5147) through a patch pipette with a manually broken tip which was then also used to remove loosened debris (*41*). Before recording, the specimen was washed with saline for

5 minutes through a gravitation perfusion system at ~ 2 ml/min, the bath volume was ~ 300 µl. Recordings were done with patch pipettes (borosilicate glass capillaries, o.d. 1.5 mm, i.d. 1 mm, without filament, World Precision Instruments, Cat# PG52151-4) pulled with a PC-10 vertical puller (Narishige, Japan) filled with internal patch solution (in mM: Kgluconate 140, Mg-ATP 2 $MgCl_2$ 2, EGTA 11, HEPES 10. pH was adjusted to 7.24 with 1 N KOH, osmolality was adjusted to 300 mOsM/kg with glucose if necessary) that approached the two MNs from opposite sides. Patch pipette tip resistance was between 5 and 6 MΩ with these solutions. Double patch recordings were performed by connecting each of the two patch pipettes to a separate Axopatch 200B patch clamp amplifier (Molecular Devices). Data were filtered at 5 kHz through a lowpass Bessel filter, digitized through an analog/digital converter (Digidata 1440), and signals were recorded with pClamp10.7 software (both Molecular Devices). Output gain was 10x. For double recordings, one MN was approached with a patch pipette. After giga seal formation, pipette capacitance artifacts were canceled manually and whole cell configuration was established at a holding potential of -70 mV. After setting whole cell capacitance compensation, correction and prediction values as well as series resistance compensation using the respective dials of the amplifier (only used to judge on recording quality as all of these compensations are disabled in current clamp mode), the recording was disconnected from external influence by using the I=0 setting of the amplifier, making it possible to revert to bath mode without disturbing the already established recording of the first MN. Now recording of the second MN was established identically. To monitor both recordings, separate channels of the digitizer and in the pClamp10.7 software were used. Quality parameters were: giga seal > 5 GΩ, membrane potential of -70 mV was held with a holding current smaller than ± 100 pA, series resistances > 15 MΩ were not accepted to ensure good control when applying current injections – only if these criteria were met for both MNs, the recordings were switched to current clamp mode. Resting membrane potential was ~ -60 mV without current injection. To determine coupling strength and rectification parameters of gap junctions between DLM-MNs, depolarizing and hyperpolarizing current was injected in one MN while the other was monitored simultaneously, and *vice versa*. Slow tonic firing was induced in one or both MNs at rates between 3-8 Hz, as is observed during flight behavior, by small somatic current injections while monitoring the respective other MN. For input-output relationships, firing was induced by 1000 ms square pulse current injections up to 1 nA in 0.1 nA increments.

<u>Intracellular dye filling</u>

For intracellular dye fills, *FMRP$^{RNAi}$* targeted to DLM-MNs (*w;UAS-FMRP$^{RNAi}$/23H06-ADZ UAS-CD4-td-GFP;30A07-DBD/+*) was used as this was reported to enhance already present dye coupling through gap junctions in Drosophila (*20*). After removing legs and wings, 2-3 days old adult male Drosophila were pinned in a Sylgard coated lid of a 35 mm Petri dish and fixed dorsal side up with two minute pins, one through the head and one through the abdomen. After submerging the specimen in normal saline (in mM: NaCl 128, KCl 2, $CaCl_2$ 1.8, $MgCl_2$ 4, HEPES 5, Sucrose ~35.5, pH was adjusted to 7.24 with 1 N NaOH, and osmolality was 300 mOsM/kg), it was opened along the dorsal midline up to the neck connectives with iris scissors. The cut dorsal longitudinal muscle (DLM) was then pinned to the sides with one minute pin each to expose gut, inner organs, and VNC. The gut, salivary glands as well as other inner organs were removed to fully expose the VNC. The specimen was then rinsed thoroughly with saline to remove excess debris. After mounting the preparation onto the stage of an upright Zeiss Axio Examiner epifluorescence microscope, the VNC was viewed with a 40x water immersion lens

(Zeiss W Plan Apochromat 40x NA 1.0, DIC VIS-R), the UAS-CD4-td-GFP-expressing DLM-MN1-5 were viewed through a FITC filter set. The ganglionic sheath was removed focally by using a broken patch pipette filled with 1% protease type XIV (from *Streptomyces griseus*, Sigma Aldrich, Cat# P5147) to allow access to DLM-MN somata (*41*). Dye fills were performed with sharp glass microelectrodes pulled from filamented borosilicate glass capillaries (Sutter BF100-50-10, tip resistance ~ 40 MΩ) with a Sutter P-97 Flaming Brown microelectrode puller. The tip was filled with a 50/50 mixture of TRITC-Dextran 3000 lysin fixable (Invitrogen, Cat# 3308) and Neurobiotin (Vector Labs, Cat# SP-1120) dissolved in 2 M KAcetate, and the shaft was then filled with 2 M KAcetate leaving an air bubble between the dye-filled tip and the shaft to avoid dilution of the dye. MNs were impaled by a short buzz (which makes the electrode tip vibrate for a set amount of time, here ~ 40 ms) with a remote buzz connected to an Axoclamp 2B intracellular amplifier (Molecular Device) that was also used for dye filling. MNs were filled iontophoretically by positive current injection (between 0.5 and 1 nA) in bridge mode until the MN was judged filled, after ~ 10 min. Then the microelectrode was removed, and the specimen was fixated with 4% paraformaldehyde at room temperature (RT) for 50 minutes, not shaking. This was followed by three rinses with PBS and 3 x 20 min PBS and 6 x 20 min washes with 0.5 % PBS-TritonX-100 (PBT, TritonX-100 Sigma Aldrich, Cat# T8787), shaking. The preparation was then incubated with Streptavidin coupled to Alexa 647 (Thermo Fisher Scientific, Cat# S-21374) in 0.3 % PBT at RT in the dark for 2 hours, shaking. Streptavidin Alexa 647 was then removed, the preparation was rinsed several times with PBS, and then washed 3 x 20 min with PBS at RT in the dark, shaking. This was followed by an ascending ethanol series 50%, 70%, 90%, 100%, 10 min each. The preparation was then mounted in methylsalicylate, topped with a high precision (170 ± 5 µm) cover slip and sealed with clear nail polish. The dye fill was then visualized with a Leica TCS SP8 confocal laser microscope with a Helium-Neon laser. Alexa 647 was excited at 633 nm and emission was detected between 650 and 680 nm with a photomultiplier tube. Images were taken with a 40x oil objective (NA 1.3) with a 1.75 digital zoom at a resolution of 1024x1024 pixels, z-step size was 1 µm.

<u>*Software*</u>

Figures were created in Corel DRAW 2021 and Adobe Illustrator 2021.

**Supplementary Text**

Two conductance based motoneuron model and gap junctional circuit model

The motoneurons are described by a single compartment model with fast spike generating $Na^+$ and $K^+$ currents, $I_{Na}$ and $I_K$ based on (*30*). The coupling is mediated by linear non-rectifying gap junction current, $I_{gap}$. The current-balance equation reads,

$$C_m \dot{v}_i = I_{in} - I_L(v_i) - I_{sb}(v_i) - I_{Na}(v_i) + \sum_{j \neq i} I_{gap}^{ij}(v_i, v_j).$$

The ionic transmembrane currents are defined as

$$I_L(v) = g_L(v - E_L),$$

$$I_{sb}(v) = g_{sb} b^4 (v - E_K),$$

$$I_{Na}(v) = g_{Na} m_\infty^3(v)(1 - h)(v - E_{Na}).$$

Gates of the voltage dependent ion channels follow first-order kinetics

$$\tau_b(v)\dot{b} = b_\infty(v) - b,$$

$$\tau_h(v)\dot{h} = h_\infty(v) - h,$$

The bi-directional, non-rectifying gap junction with linear charge transfer (Figs. 2E, F) are described by the following current

(1) $I_{gap}^{ij}(v_i, v_j) = g_{gap}^{ij}(v_i - v_j).$

Parameters common to all simulations are found in Table S2. For the simulations with heterogeneous coupling (Fig. 3G), coupling strengths were chosen to match the coupling coefficients measured experimentally in Fig. 2F. This yielded $g_{gap}^{12} = g_{gap}^{21} = g_{gap}^{34} = g_{gap}^{43} = 203.32pS$ and 90.37 pS for the remaining connections. To keep cases as comparable as possible, the coupling strength for homogeneous coupling (Fig. 3H) was chosen to be the mean of the connections in the heterogenous case with $g_{gap}^{ij} = 128.02pS \forall i \neq j$. The same coupling strength was used for Fig 3D, except for in the strong gap junction HOM case, where the coupling strength was 3nS.

Stochastic and deterministic simulations of the model are performed using the `brian2` package for python (*42*). The stochastic Heun method was used with a time step of 3us. For noisy simulations, a zero-mean white-noise current was added to the current balance equation (using the brian2 `xi` variable) with a noise strength of σ = $1.5811pA\sqrt{ms}$ (Fig. 3D, G and H) or σ = $0.1pA\sqrt{ms}$ (Fig. 3C).

In addition, $Q$ represents $\frac{e}{k_B T}$, which is the elementary charge $e$ divided by the Boltzmann constant $k_B$ and the temperature $T$ in Kelvin. Throughout the simulations, $Q$ is approximated as $39.2/V$.

The model parameters common to all models are summarized in Table S2, while the activation curves of the gates and the kinetic time scales are provided in Table S3. Some parameters and equations were reported inconsistently in the original publication. Thanks to the authors kindly providing their code, we were able to extract the parameters used for their simulations, which are reported here. Additionally, to produce different onset bifurcations, the shab conductance was changed (see below) and the input current was adjusted to move the system close to onset. The values differing between models are summarized in S4.

For all simulations, neurons were initiated in random phases, by drawing parameter combinations from random time points of simulations of the same neuron model periodically spiking without coupling.

Gap junctions, of the form Eq. (1), influence the intrinsic properties of neurons, since they act as effective leak and capacitance changes. Hence $g_{gap}$ can even induce bifurcations in the single neuron (*29*). In Fig. 3C the coupling is homogenous $g_{gap}^{i,j} = g_{gap} \; \forall i \neq j$ between pairs but coupling strength $g_{gap}$ is varied. Here, the passive properties of the membrane were compensated via $g_L^{new} = g_L - (N-1)g_{gap}$ and $I_{in}^{new} = I_{in} + (N-1)g_{gap}50mV$. This stabilizes firing rates and PRC shape while changing $g_{gap}$.

Shab-induced bifurcations

A local bifurcation analysis of the fixpoints in the model was shown in (*30*). The important quantity for predicting network states of the CPG is the phase response curve (Fig. 3E), which is closely related to the spike-onset bifurcation of the neuron, *i.e.*, the excitability class. The bifurcation diagram in Fig. S7A adds nonlocal codim-2 bifurcations that organize transitions between excitability classes and their PRCs (labeled SNL, NSL, FLC/Hopf)(*43*). Depending on the shab channel density, $g_{sb}$ on the ordinate, different onset bifurcations that change the resting state into the spiking mode occur. In the middle range of $g_{sb}$ levels, spiking commences via a well-known saddle-node on invariant cycle (SNIC) bifurcation. The SNIC region is encapsulated by two saddle-node loop points (SNL). At the lower end, for $g_{sb}$ smaller than the small saddle node loop (sSNL) point, spiking commences via a small homoclinic loop (HOM, green line) at lower input currents, $I_{in}$ than the saddle-node bifurcation (blue line). At the upper end of the SNIC range, a series of bifurcations is traversed that eventually leads to spike onset via a fold of limit cycles (FLC) and a subsequent subcritical Hopf bifurcation, termed Hopf regime here for simplicity. Bifurcations leading into that excitability class are the big saddle-node loop (bSNL), the neutral saddle loop (NSL) and also a cusp (CP). The Bogdanov-Takens point in between creates a Hopf line (AH, red line) that eventually turns back to create the excitation block at higher input levels.
The consequences for the phase relations of coupled neurons can be understood from the neuron's phase response curves (PRC) at different bifurcations (Fig. S7A insets). Just as the ability to produce arbitrarily low firing rates, PRC shapes are associated with the neuron's excitation class (onset bifurcation) (*44-46*). The insets in Fig. S7B show prototypical PRCs for the three onset bifurcations, which can be achieved by different Shab levels. The exact parameters are summarized in Table S2. A critical transition with repercussions for the CPG network states is the symmetry breaking in the PRC at the sSNL bifurcation as detailed in the next section of the supplement.

A FLC/Hopf onset bifurcation as spike onset can be ruled out as a model for the biological MN1-5 network based on the following grounds: The region where tonic spiking commences via the FLC/Hopf bifurcation has a finite, nonzero frequency at the rheobase. For the present model (upper arrow in Fig. S7C) this is at around 47 Hz and hence above the dynamic range of the *in vivo* measures $f$-$I$ curves from the motoneurons, which operate between 0-40 Hz (3-25 Hz during normal flight), see Fig. 1E.

The bifurcation analysis was carried out using `AUTO-07P` (*47*).

Phase-reduced circuit model with repulsive coupling

Since the cholinergic inputs drive MN1-5 into a tonically spiking regime, it is possible to reduce the conductance-based neurons to approximate phase-oscillators (*31*). The $i^{\text{th}}$ phase oscillator is defined by its intrinsic, mean firing rate, $f_i$, and its phase response curve, $Z(\phi_i)$, obtained from the biophysical model (Fig. S7A). The PRC are calculated based on a direct perturbation approach (*48,49*). The network equation reads,

(2) $\dot{\phi}_i = f_i + \sum_{k \neq i} Z(\phi_i) g(\phi_i, \phi_k)$,

where $g(\phi_i, \phi_j)$ is the phase dependent perturbation received via the gap junctions. It depends on the phases of the pre- and post-synaptic neurons. If the spike waveforms are idiosyncratic, the interaction term, based on Eq. (1) reads,

$$g(\phi_i, \phi_k) = g_{\text{gap}}(v(\phi_k) - v(\phi_i))/C_{\text{m}}.$$

To understand the influence of the PRC on phase relations in small networks, first consider the phase difference between two coupled neurons $i$ and $j$, isolated from the network. Define their phase difference $\psi = \phi_i - \phi_j$. Then, using approximate averaging theory, the slow evolution of the phase difference simplifies to (*28*),

(3) $\dot{\psi} = \nu + G^{odd}(\psi)$.

Here, the small frequency detuning is $\nu = f_i - f_j$ and the averaged coupling function is defined as

(4) $G(\psi) = \int_0^1 Z(\phi_i) g(\phi_i, \phi_i + \psi) d\phi_i$.

Stable fixpoints of Eq. (3) determine constant phase relations for neurons with similar intrinsic frequencies $\nu$ as illustrated in Figure S7. In Fig. 3E the odd part $G^{odd}(\psi) = G(\phi) - G(-\phi)$ and the fixpoints are shown for different choices of $Z(\phi)$. The stable in-phase fixpoint directly translates to a synchronous network state also for network sizes of $N > 2$. This is, however, not observed in the CPG recordings, Fig. 1A.

For the network to show frustration, pairs of neurons need to have a stable phase fixpoint in antiphase, $\psi = \phi_i - \phi_j = 0.5$, *i.e.*, they must be phase-repellent.

The top spike trains in Fig. S7B show an example based on a HOM PRC that generates stable antiphase fixpoints. The stability can be analyzed using the iterated map of the phase differences. The return map reads $\psi_{n+1} = \psi_n + G^{\text{odd}}(\psi_n)$. Fig. S7C illustrates how asymmetric PRCs lead to a stable fixpoint in antiphase (left). The SNIC case is shown in Fig. S7B, bottom, and Fig. S7C, left, and shows in-phase synchronization.

In conclusion, reciprocally coupled neurons, like those connected via non-rectifying gap junctions, require an asymmetric coupling function to show stable phase locking. This can be achieved via an asymmetric PRC. For the antiphase to be stable and the network in Eq. (2) to show frustration, the coupling function has to show phase advance in the first half of the ISI and phase delay in the latter half. This property is found in the HOM PRC. This class of PRCs is only found for $g_{sb}$ conductances below the sSNL bifurcation (Fig. S7A). Fig. 3E shows that out of the three spike-onset bifurcations found in the neuron model, the HOM regime is the only one exhibiting a stable antiphase fixpoint leading to a splay state in the frustrated network.

The splay state measure

In order to quantify the splayness of a network, statistics of phase-differences are calculated. In particular, two measures are applied. First, phase-differences of neurons are compared to those that would arise from a perfect splay state. The phases are interpolated between spikes as

$$\phi_i(t) = \frac{t - t_{n,i}}{t_{n+1,i} - t_{n,i}}, (t_{n,i} \leq t < t_{n+1,i})$$

Then for each point in time, the phases are ordered such that

$$\phi_k(t) > \phi_l(t), (k > l)$$

The phase differences are computed as

$$\psi_i(t) = \phi_{i+1}(t) - \phi_i(t)\ \forall i = 0,1, \dots N - 1$$

and

$$\psi_N(t) = 1 - \sum_{i=1}^{N} \psi_i\,(t)$$

In order to calculate the splayness, these phase differences are compared to those of the most splayed state (splay state) and that of the least splayed state (sync state):

(5) $r(t) = \frac{\sum_i^N (\psi_{i,t} - \frac{1}{N})^2}{(N-1)N^{-2} + (1 - N^{-1})^2}$

From this we construct the instantaneous splayness

(6) $s(t) = 1 - \sqrt{r(t)}$

And a time-averaged splayness index

(7) $s = 1 - \sqrt{\frac{1}{T}\sum_t r(t)}$

Second, classical order parameters are evaluated

$r_k = \sum_{j=1}^{N} e^{i2\pi k \phi_j(t)}$.

The time-averaged $\langle r_4 \rangle$ indicates splayed activity, while the time-averaged $\langle r_1 \rangle$ measures in-phase synchrony.

Lastly, a spectral clustering of the synchronization matrix $m_{jk} = \langle e^{i(\phi_j(t) - \phi_k(t))} \rangle$ was used to visually confirm that indeed the order parameter correspond to splay and sync states.

The code for the computational models will be made available upon request.

Supplementary figures

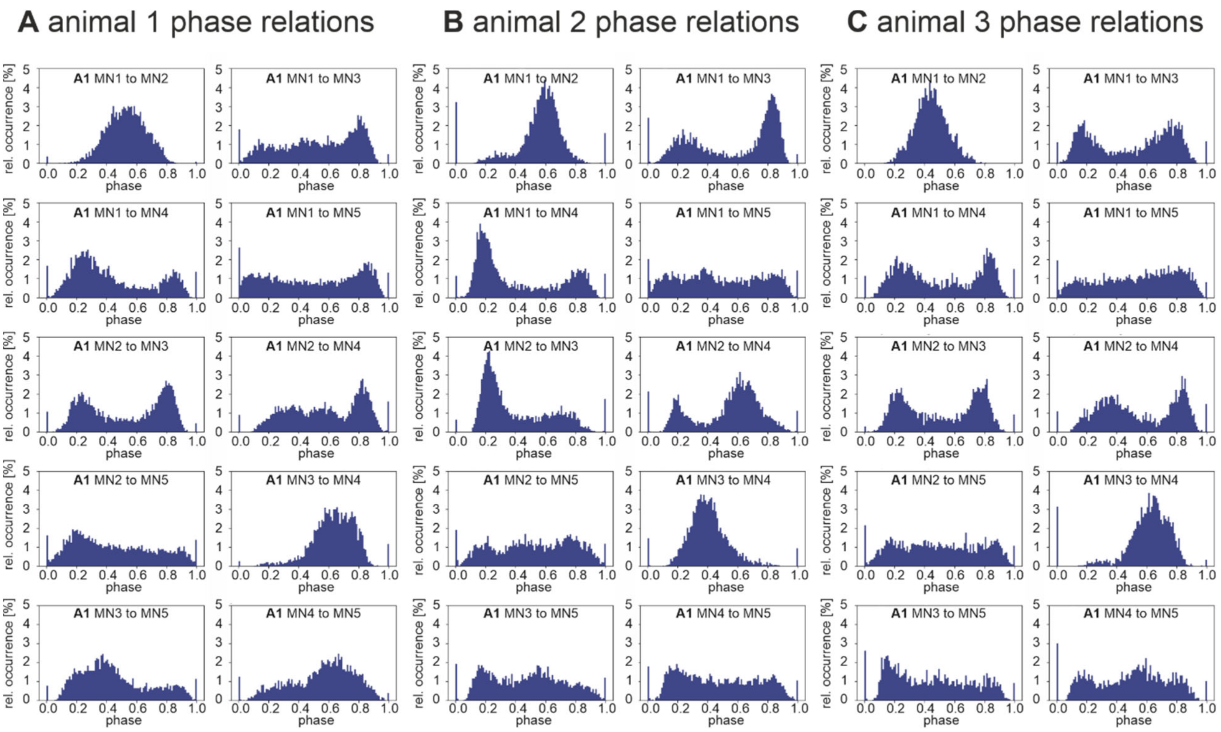

**Fig. S1.**
*Firing phase relationships of DLM-MN pairs are conserved between individuals.*
For three individual male flies (A-C) all 10 possible pairwise combinations of the 5 DLM-MNs (MN1-5) are plotted as cumulative phase histograms from simultaneous recordings of MN1-5 during 10 minutes of continuous tethered flight (equaling approximately 3000 spikes of each MN). Starting with the MN1/MN2 pair on the upper left, (A1-C1), for each MN pair, the interspike intervals between two consecutive spikes of one MN were divided into 100 equally sized bins (x-axis), and it was determined in which bin the spike of the other MN occurred. This was repeated as sliding window for all interspike intervals to fill the bins cumulatively. Cumulative spike counts in each bin were normalized to total spike count (y-axis). In all three individuals (A-C) different DLM-MN pairs show different phase relationships, but the same pairs show similar phase relationships across individuals.

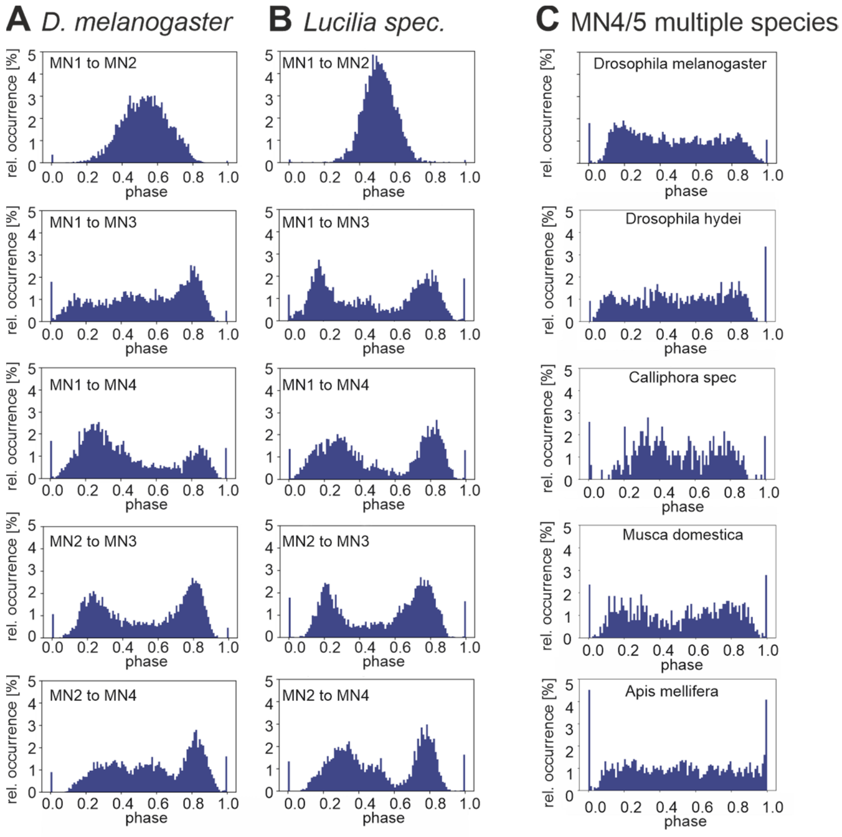

**Fig. S2.**
*Phase relations between DLM-MNs are conserved across species.*
In all tested flying insect species, the dorsal longitudinal wing depressor muscles consists of 6 muscle fibers, each of which is innervated by one MN, namely MN1-5 (for schematic see Fig. 1). In *Drosophila melanogaster* each of the 10 MN1-5 pairs shows characteristic phase relationships in their firing patterns (see Figs. 1 and Fig. S1). Analyses of *in vivo* recordings during tethered flight reveals highly similar firing phase relationships of different DLM-MN pairs in *Drosophila melanogaster* (A) and in the goldfly, *Lucilia spec* (B), a related dipteran species. (C) *In vivo* recordings of the MN4/MN5 pair shows similar firing phase relationships between the fruit fly *Drosophila melanogaster* (top), another Drosophilidae (*Drosophila hydei*, second from top 3), two additional dipteran species (the blowfly *Calliphora spec.*, third from top; and the house fly, *Musca domestica*, second from bottom). Some characteristics of the phase relationships observed in all dipteran species tested (top 4 panels) are also observed in a hymenopteran species, the honey bee *Apis melifera* (bottom). There the characteristic inhibition of MN4 firing just after MN5 spikes is recapitulated, but the depression of MN4 firing just before MN5 spikes is not.

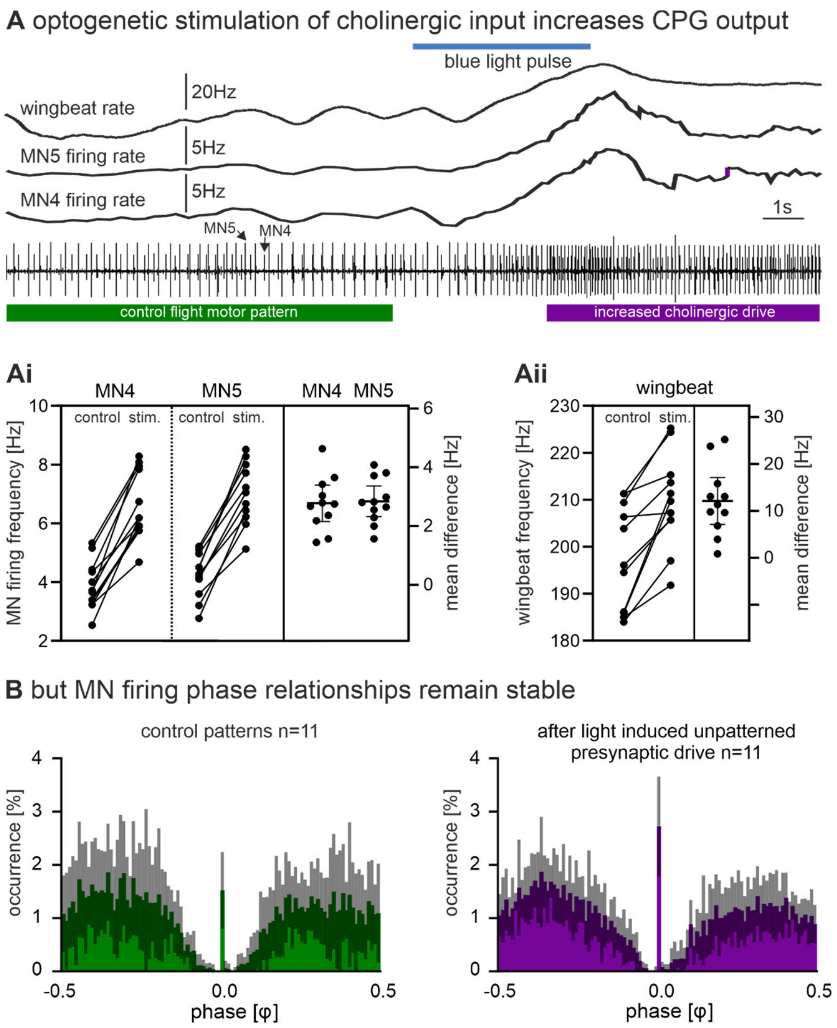

**Fig. S3.**

*Unpatterned excitatory input scales up CPG output without affecting the pattern.*

(A) The bottom trace shows a representative recording of MN4 (arrow, smaller unit) and MN5 (arrow, large unit) with one tungsten electrode during tethered flight. Timing of unpatterned optogenetic activation of presynaptic cholinergic interneurons is indicated by a blue line. Upper three traces show the instantaneous wingbeat frequency (top trace) and the instantaneous firing frequencies of MN5 and MN4. (Ai) Quantification from 11 animals shows that optogenetic activation of cholinergic interneurons increases the firing frequencies of both MNs significantly and to the same degree. (Aii) Consequently, wingbeat frequency is increased significantly. (B) The typical phase relationship of MN4 and MN5 firing that is observed in controls (Figs. 2A, S1) and prior to optogenetic stimulation (left) remains unaltered upon increasing CPG output by unpatterned stimulation of cholinergic interneurons (right).

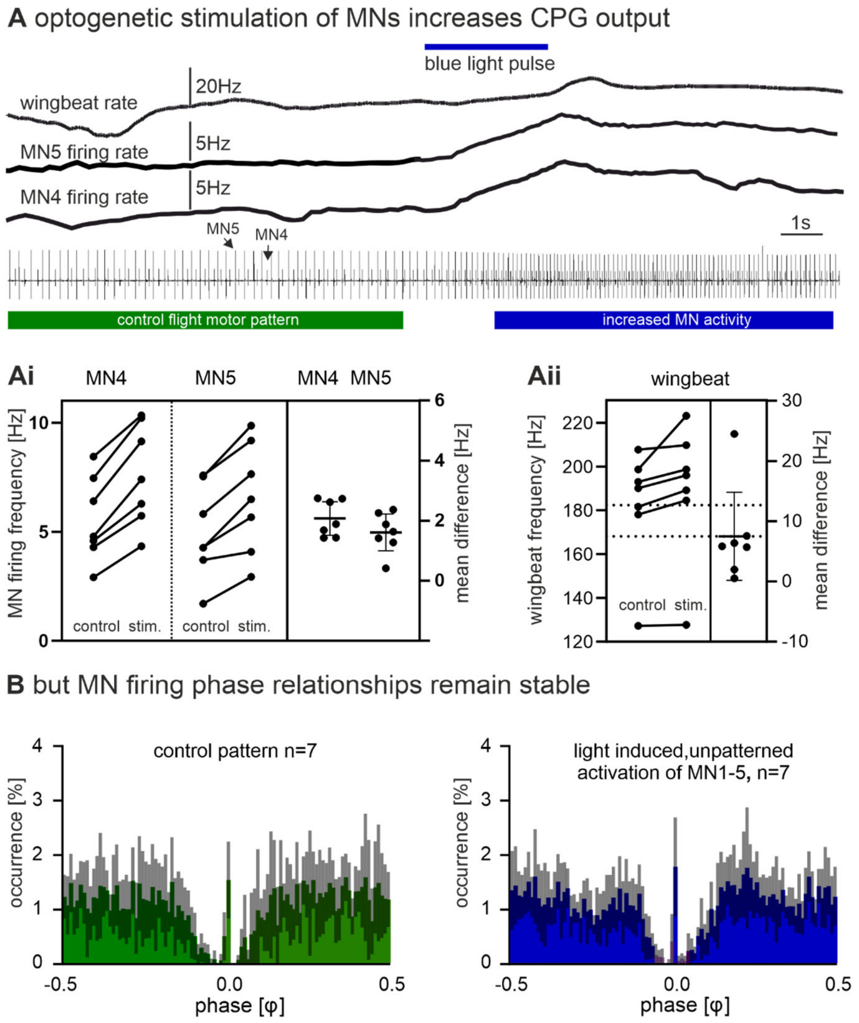

**Fig. S4.**
*Unpatterned optogenetic activation of DLM-MNs during flight scales up CPG output without affecting the pattern.*
(A) The bottom trace shows a representative recording of MN4 (arrow, smaller unit) and MN5 (arrow, large unit) with one tungsten electrode during tethered flight. Timing of unpatterned optogenetic activation DLM-MNs is indicated by a blue line. Upper three traces show the instantaneous wingbeat frequency (top trace) and the instantaneous firing frequencies of MN5 and MN4. (Ai) Quantification from 7 animals shows that optogenetic activation increases the firing frequencies of both MNs significantly and to the same degree. (Aii) Consequently, wingbeat frequency is increased significantly. (B) The typical phase relationship of MN4 and MN5 firing that is observed in controls (Figs. 2A, S1) and prior to optogenetic stimulation (left) remains unaltered upon increasing CPG output by unpatterned stimulation the DLM-MNs (right).

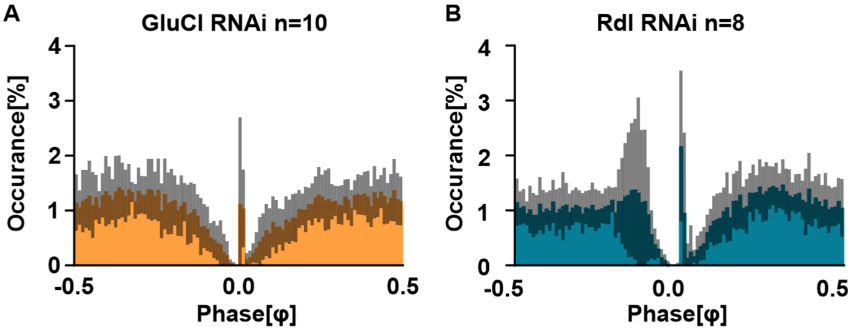

**Fig. S5.**
*Inhibitory chemical synapses do not affect MN phase relationships during flight.*
Firing phase relationships between MN4 and MN5 as observed in control animals (Figs. 2A, Fig. S1) remain unaltered upon targeted RNAi knockdown of receptors at the two predominant inhibitory chemical synapses to DLM-MNs, (A) the glutamate gated chloride channel (GluCl) and (B) Rdl GABA-ARs. GluCl-RNAi knockdown efficacy was confirmed by Western blotting (Ai) and Rdl GABA-AR knockdown efficacy has previously been confirmed (Ryglewski et al., 2017).

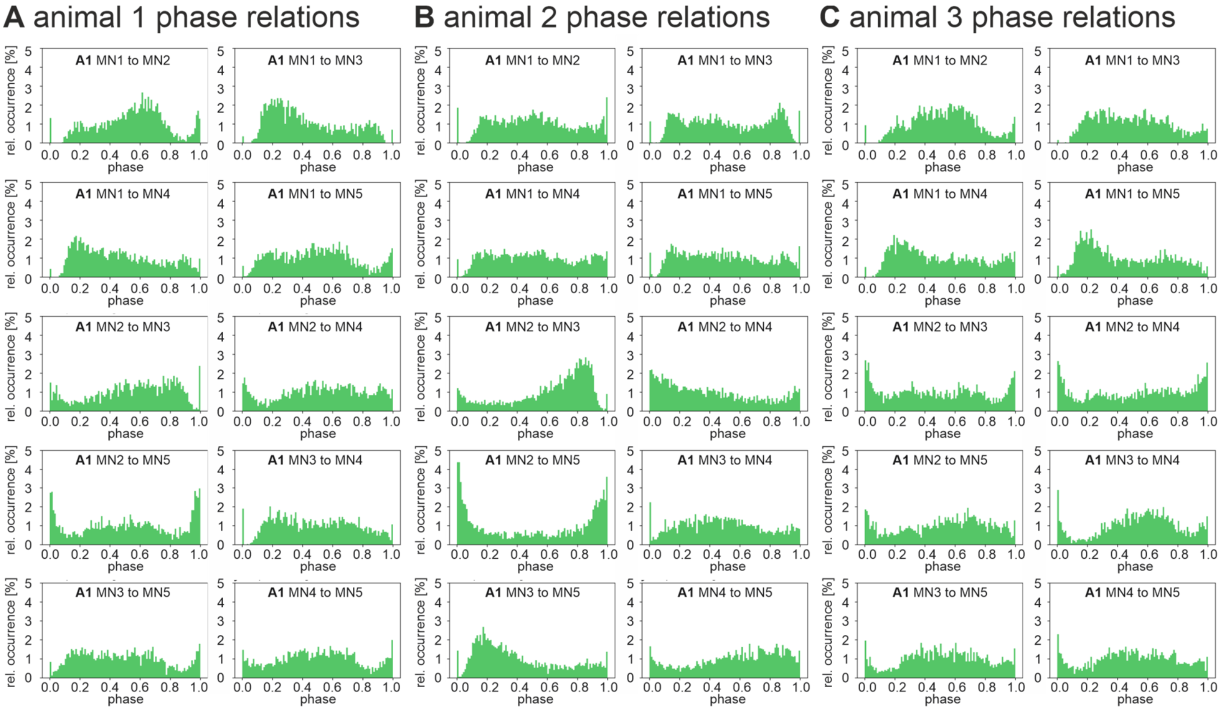

**Fig. S6.**

*DLM-MN firing phase relationships require GJs between MNs.*

For three individual male *Drosophila melanogaster* flies with targeted expression of RNAi knockdown in MN1-5 (A-C) all 10 possible pairwise combinations of the 5 DLM-MNs (MN1-5) are plotted as cumulative phase histograms from simultaneous recordings of MN1-5 during 10 minutes of continuous tethered flight (equaling approximately 3000 spikes of each MN). Starting with the MN1/MN2 pair on the upper left, (A1-C1), for each MN pair, the interspike intervals between two consecutive MN spikes was normalized and divided into 100 equally sized bins (x-axis). Next it was determined in which bin the spike of the other MN occurred. This was repeated as sliding window for all interspike intervals and bins were filled cumulatively and normalized to total spike count (y-axis). In all three individuals with knockdown for ShakB mediated GJs in MN1-5 (A-C) the phase relationships that are characteristic for each DLM-MN pair in control flies (see Fig. S1) and conserved across dipteran species (see Fig. S2) are impaired.

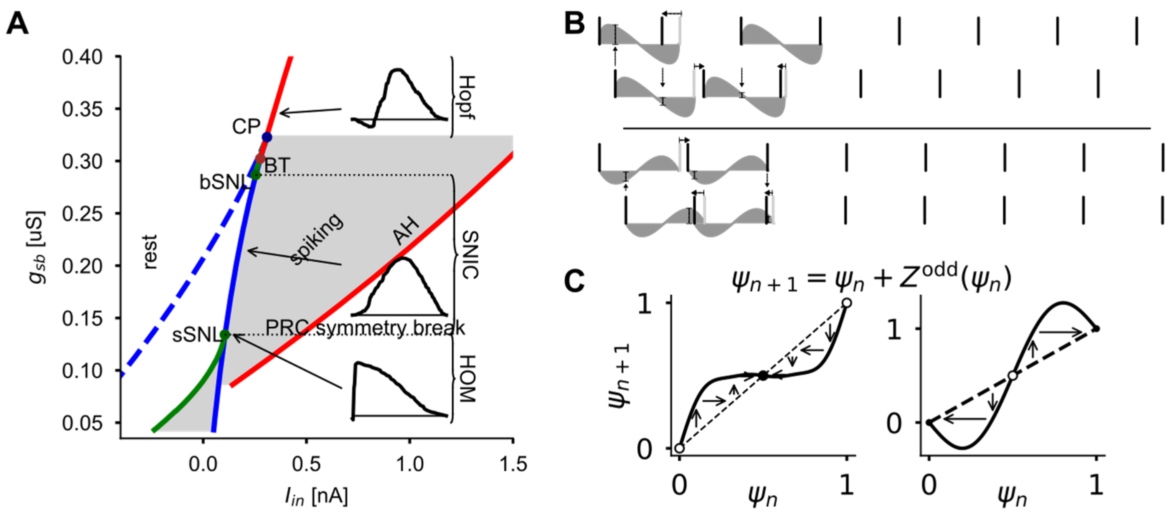

***Fig. S7.***
*Motoneuron model analysis.*
(A) Codim-2 bifurcation diagram of the MN model with focus on global bifurcations that switch the PRC shape. Small and big saddle-node loop (sSNS, bSNL) encapsulate the SNIC interval, where spiking (grey area) commences after a saddle-node bifurcation (blue line) on a limit cycle. Saddle Homoclinic loop (HOM) line is depicted in green. The red line shows the Andronov-Hopf bifurcation (AH), which ends in a Bogdanov-Takens point. At the cusp (CP) the neuron model switches from 3 to 1 fixpoint. The three PRCs in the insets show distinct symmetry properties. (B) Asymmetric PRC can lead to stable phase relations. PRCs with phase-advance after in the beginning of the ISI and phase-delay in the second half leads to anti-phase synchronization (top), while the reverse leads to in-phase synchronization (bottom). The grey areas between spikes indicate the coupling function, i.e., the phase shifted created by perturbation at that time point. (C) Iterated maps for the phase difference between two coupled neurons. The HOM case on the left and the SNIC case on the right.

**Table S1:** Reagents.

| REAGENT or RESOURCE | SOURCE | IDENTIFIER | |
|---|---|---|---|
| Chemicals, Peptides, and Recombinant Proteins | | | |
| Streptavidin Alexa 647 | Thermo Fisher Scientific | Cat# S-21374 | |
| TRITC-Dextran 3000 lysin fixable | Invitrogen | Cat# 3308 | |
| Neurobiotin Tracer | Vector Labs | Cat# SP-1120 | |
| TritonX-100 | Sigma Aldrich | Cat# T8787; CAS 9002-93-1 | |
| Protease Type XIV from *Streptomyces griseus* | Sigma Aldrich | Cat# P5147; CAS 9036-06-0 | |
| Sodium nitrite | Sigma Aldrich | Cat# 563218; CAS 7632-00-0 | |
| Potassium hydroxide | Sigma Aldrich | Cat# 306568, CAS 1310-58-3 | |
| Methylsalicylate | Sigma Aldrich | Cat# M6752; CAS 119-36-8 | |
| Sylgard 184 Dow Corning | Biesterfeld Spezialchemie Hamburg | Sylgard 184 | |
| Experimental Models: Organisms/Strains | | | used in Figure |
| *D. melanogaster*: $ShakB^{RNAi}$ in DLM MNs<br>$\frac{w}{\neg};\frac{GMR23H06-pBPp65ADZpUw\}attp40}{+};\frac{GMR30A07-pBPZpGAL4.BD.Uw\}attP2}{P\{TRiP.HMC04895\}attP2}$ | the GMR23H06-ADZ;30A07-DBD DLM Split GAL4 stock was made in our own lab with plasmids as described in the HHMI Janelia Farm Research Campus Fly Light Collection by G. Rubin (Jenett et al. 2012) | RRID: BDSC_57706 | Fig.2<br>Fig.S6 |
| *D. melanogaster*: $\mathrm{ShakB}^{OE}$ in DLM MNs<br>$\frac{w}{\neg};\frac{GMR23H06-pBPp65ADZpUw\}attp40}{UAS-shakb(N+16)};\frac{GMR30A07-pBPZpGAL4.BD.Uw\}attP2}{+}$ | DLM Split GAL4 | RRID: BDSC_86262 | Fig.2<br>Fig.4 |
| *D. melanogaster*: $\mathrm{ShakB}^{KO}$ in DLM MNs<br>$\frac{w[1118]}{\neg};\frac{P\{TKO.GS01152\}attP40}{+};\frac{P\{UAS-Cas9.C\}attP2B}{P\{GMR23H06-GAL4\}attP2}$ | | RRID:<br>BDSC_78593<br>BDSC_54595<br>BDSC_49050 | |
| *D. melanogaster*: control for *RNAi* in DLM MNs<br>$\frac{w}{\neg};\frac{GMR23H06-pBPp65ADZpUw\}attp40}{+};\frac{GMR30A07-pBPZpGAL4.BD.Uw\}attP2}{P\{UAS-GFP.VALIUM10\}attP2}$ | DLM Split GAL4 | RRID: BDSC_35786 | Fig.1<br>Fig.2 |
| *D. melanogaster*: $GluCl\alpha^{RNAi}$ mainly in DLM MNs<br>$\frac{w[1118]}{\neg};;\frac{P\{KK109167\}VIE-260B}{P\{GMR23H06-GAL4\}attP2}$ | | RRID:<br>FlyBase_FBst0477580<br>BDSC_49050 | Fig.S5 |

| | | | |
|---|---|---|---|
| *D. melanogaster*: *Rdl*$^{RNAi}$ mainly in DLM MNs<br>$\frac{w[1118]}{\neg};;\frac{P\{GD4609\}v41103}{P\{GMR23H06-GAL4\}attP2}$ | | RRID:<br>FlyBase_FBst0463935<br>BDSC_49050 | Fig.S5 |
| *D. melanogaster*: Channelrhodopsin XXL in DLM MNs<br>$\frac{w[1118]}{\neg};\frac{GMR23H06-pBPp65ADZpUw\}attp40}{P\{UAS-ChR2.XXL\}VK00018};\frac{GMR30A07-pBPZpGAL4.BD.Uw\}attP}{+}$ | DLM Split GAL4 | RRID:<br>BDSC_58374 | Fig.S4 |
| *D. melanogaster*: Channelrhodopsin XXL in cholinergic neurons<br>$\frac{w[1118]}{\neg};\frac{P\{UAS-ChR2.XXL\}VK00018}{P\{w[+mC]=ChAT-GAL4.7.4\}19B}$ | | RRID:<br>BDSC_6798<br>BDSC_58374 | Fig.S3 |
| *D. melanogaster*: *FMRP*$^{RNAi}$ in GFP-expressing DLM MNs<br>$\frac{w}{\neg};\frac{P\{GMR23H06-ADZ\}attp40\ P\{UAS-CD4-tdGFP\}8M2}{P\{KK107935\}VIE-260B};\frac{P\{GMR30A07-DBD\}attP2}{+}$ | DLM Split GAL4 | RRID:<br>BDSC_35839<br>FlyBase_FBst0482375 | Fig.2 |
| *D. melanogaster*: fast GCaMP7f expression in DLM muscle<br>$\frac{w}{\neg};\frac{P\{20XUAS-IVS-jGCaMP7f\}su(Hw)attP5}{+};\frac{GMRP\{Act88F-GAL4.1.3\}3}{+}$ | Sun, Y., Kim, D. (2019.1.30). UAS-GCaMP7 constructs and insertions from Yi Sun and Douglas Kim. Fly Base ID FBrf0241300 | RRID:<br>BDSC_38461<br>BDSC_80906 | Fig.4 |
| Software and Algorithms | | | |
| Amira 4.1 with custom plug-ins | FEI Hillsboro, Oregon, US; Schmitt el al. 2004, Evers et al. 2005 | Amira 3D analysis , RRID:SCR_014305 | |
| Corel Draw 2021 | Corel Corporation | | |
| SPSS Statistics 22 | IBM | SPSS, RRID:SCR_002865 | |
| Huygens Professional software (version 19.10.0) | Scientific Volume Imaging | RRID:SCR_014237 | |
| IR illuminator with 48 LEDs | Sygonix | | |
| pClamp10.7 electrophysiology software | Molecular Devices | pClamp {SCR:011323} | |
| Graphpad Prism 9.2.0 | GraphPad Software, San Diego, USA | RRID:SCR_002798 | |
| Spike2 (Version 7.2) | CED | | |
| FASTCAM Viewer software (PFV Version 3.6.9.0) | Photron | | |
| Leica Application Suite X (LASX) | Leica Microsystems | | |
| Illustrator 2021 version 25.2 | Adobe | | |
| Jupyter Notebook (custom Python routines) | | | |
| Python library NumPy | | | |
| Python library pickle | | | |
| Python library SciPy | | | |
| Python library Matplotlib | | | |
| Other | | | |
| Glass microelectrodes with filament, i.d. 0.5 mm, o.d. 1.0 mm | Sutter Instruments | Cat# BF 100-50-10 | |

| | | |
|---|---|---|
| Dental Cromalux-E Halogen Curing Light Unit | Mega Physik | REF 7050.B01 |
| clear glass adhesive | Duro; Pacer Technology, Rancho Cucamonga, CA | |
| Grass SD9 square pulse stimulator | | |
| Sygonix IR illuminator with 48 LEDs | | |
| Axopatch 200B Patch Clamp amplifier | Molecular Devices | |
| Digidata 1440 or 1550B analog/digital converter | Molecular Devices | |
| Patch glass without filament, i.d. 1.0 mm, o.d. 1.5 mm | World precision instruments | Cat# PG52151-4 |
| PC-10 vertical pipette puller | Narishige | |
| P-97 Flaming Brown glass microelectrode puller | Sutter | |
| Osmomat 3000 basic freezing point osmometer | Gonotec | |
| Drosophila food vials, polystyrene, 68 ml | Kisker | Cat# 789008 |
| Model 1700 extracellular amplifier | AM-Systems | |
| FASTCAM Mini UX100 | Photron | |
| tungsten rods, diameter 127 μm | Science products | Cat# TW5-4 |
| tungsten wire, diameter 100 μm | MaTeck | Cat# 009486-1 |
| AxioExaminer A1 with Zeiss W Plan Apochromat 40x NA 1.0, DIC VIS-R lens | Zeiss, Germany | |
| TCS SP8 laser scanning confocal microscope with 40x oil objective | Leica Microsystems, Germany | |

**Table S2:** Model parameters common to all models.

| Parameter | Value |
|---|---|
| $C_m$ | 0.13 nF |
| $g_{Na}$ | 0.4312 uS |
| $g_L$ | 0.008624 uS |
| $E_L$ | -60 mV |
| $E_K$ | -72 mV |
| $E_{Na}$ | 55 mV |
| $z_m$ | 3 |
| $v_m$ | -33 mV |
| $z_h$ | 5.2 |
| $v_h$ | -39.14 mV |
| $r_h$ | 0.2/ms |
| $\gamma_h$ | 0.38 |
| $z_b$ | 1.1056 |
| $v_b$ | -42.14 mV |
| $r_b$ | 0.2/ms |
| $\gamma_b$ | 0.38 |

**Table S3:** Activation functions and kinetic time scales of different gates.

| gate | activation curve / kinetic timescale |
|---|---|
| $p_\infty$ | $\frac{1}{1+e^{-Qz_p(v-v_p)}}$ |
| $\tau_p$ | $\frac{e^{-Qz_p\gamma_p(v-v_p)}}{r_p(1+e^{-Qz_p(v-v_p)})}$ |

**Table S4:** Model parameters differing between models.

| Parameter | Hopf | SNIC | HOM/SNL |
|---|---:|---:|---:|
| $g_K$ | 0.34496 uS | 0.2156 uS | 0.13768216 uS |
| $I_{in}$ | 330 pA | 175 pA | 108.3 pA |

**Movie S1.**

High speed video (6400 FPS) of *Drosophila melanogaster* during tethered flight with 3 recording electrodes.

**Movie S2.**

High speed video with (5000 FPS) of *Lucilia spec.* during tethered flight with 2 recording electrodes.

**Movie S3.**

High speed video with (5000 FPS) of A*pis mellifera* during tethered flight.
